# Open-source LED lamp for the LI-6800 photosynthesis system

**DOI:** 10.1101/2023.10.18.562806

**Authors:** Aarón I. Vélez Ramírez, Juan de Dios Moreno, Uriel G. Pérez-Guerrero, Antonio M. Juarez, Hector Castillo-Arriaga, Josefina Vázquez-Medrano, Ilane Hernández-Morales

## Abstract

**Premise:** Controlling light flux density during carbon dioxide assimilation measurements is essential in photosynthesis research. Commercial lamps are expensive and are based on monochromatic light emitting diodes (LEDs), which deviate significantly in their spectral distribution compared to sunlight.

**Methods and Results:** Using a LED emitting white light with a color temperature similar to sunlight, we developed a cost-effective lamp compatible with the LI-6800 photosynthesis measuring system. When coupled with customized software, the lamp can be controlled via the LI-6800 console by a user or Python scripts. Testing and calibration show that the lamp meets the quality needed to estimate photosynthesis parameters.

**Conclusions:** The lamp can be built using a basic electronics lab and a 3D printer. Calibration instructions are supplied and only require equipment commonly available at plant science laboratories. The lamp is a cost-effective alternative to perform photosynthesis research coupled with the popular LI-6800 photosynthesis measuring system.

## INTRODUCTION

A common method to study photosynthesis in vivo requires measuring the net assimilation rate (A) at various environmental conditions, which are carefully set and controlled (Busch, 2018; Coe and Lin, 2018; Walker et al., 2018). While ambient CO_2_ and irradiance (I) are changed, A is measured, resulting in two dose-response curves. The first curve is known as A-Ci curves as changes in ambient CO_2_ result in changes in the intracellular CO_2_ concentration (Ci) (Busch, 2018). The second curve is known as A-I (Coe and Lin, 2018). In the International System of Units, I is measured in W·m^−2^ (J·m^−2^·s^−1^); in plant sciences, however, quantum units are preferred as A increases in response to the number of incident photons (measured in µmol) with wavelengths between 400 and 700 nm regardless of their energy content (measured in J) (Wayne, 2016; Stanghellini et al., 2019). Here, therefore, we measure the light flux density in quantum units (µmol·m^−2^·s^−1^) and referred to as photosynthetically active photon flux density (PPFD), resulting in A-PPFD curves. The measurements of one or both dose-response curves can be used to estimate photosynthetic parameters (Yin et al., 2009; Velez-Ramirez et al., 2014). The photosynthetic parameters are numeric representations of subprocesses that together contribute to A (Busch, 2018; Coe and Lin, 2018). The most common model used to calculate the photosynthetic parameters is the one proposed by Farquhar et al. (Farquhar et al., 1980).

Measuring either dose-response curves requires a way to precisely control the PPFD incident on the leaf sample (Busch, 2018; Coe and Lin, 2018). Manufacturers of photosynthesis measuring systems use lamps based on light emitting diodes (LEDs); for example, the LI-6400 and LI-6800 from Li-Cor Biosciences (Lincoln, Nebraska, USA), and the GFS-3000 from Heinz Walz (Effeltrich, Germany) have LED lamps as optional accessories. In these systems, however, color LEDs are used. For instance, the LI-6800 basic lamp (model 6800-02) uses blue and red LEDs. Growing plants under light sources with a spectral distribution deviating from solar light results in significant morphological and physiological changes (Hogewoning et al., 2010a). When using only red and blue LEDs to grow plants, some plants may show a significant decreases in photosynthesis if blue light proportion is low compared to red (Hogewoning et al., 2010b; Trouwborst et al., 2016). In addition to the light spectrum during the growth period, the spectral distribution of the light used during photosynthesis measurements affects the shape of the A-PPFD curves, resulting in different A values at the same PPFD (Liu and van Iersel, 2021). Hence, the commercial LED lamps for measuring photosynthesis have the disadvantage of supplying light from only monochromatic LEDs. Additionally, the high price for these lamps — considering that are mainly composed of LEDs and their drivers — calls for the development of an open-source lamp that uses white LED light. Here, we report the development of such a lamp to be used with the LI-6800 photosynthesis measuring system.

A mayor design goal was to make the lamp compatible with script-based measurements. That is, to fully integrate the proposed lamp with the LI-6800 background programs (BP), which are Python scripts that autonomously perform complex measurement procedures (like dose-response curves). We achieved this integration by using a custom-made LED driver, the LI-6800 *User I/O* port and an Arduino-compatible (www.arduino.cc) microcontroller board, programed with Python. Hardware, software, detailed building instructions, and calibration procedures are provided under open licenses.

## METHODS AND RESULTS

### Lamp hardware

#### COB-LED and LED driver

The Lamp described here is based on a Chip on Board-Light Emitting Diode (COB-LED) array, emitting white light with a color temperature of 6500 K (Table 1), which corresponds to the color temperature of an average daylight (International Commission on Illumination, (Judd et al., 1964)). The COB-LED array has a forward voltage of 18 V and can operate at a current up to 1400 mA. In this design, however, its maximal operating current is set at 1000 mA. Hence, the maximal power consumed by the LED is 18 W.

**Table 1.**
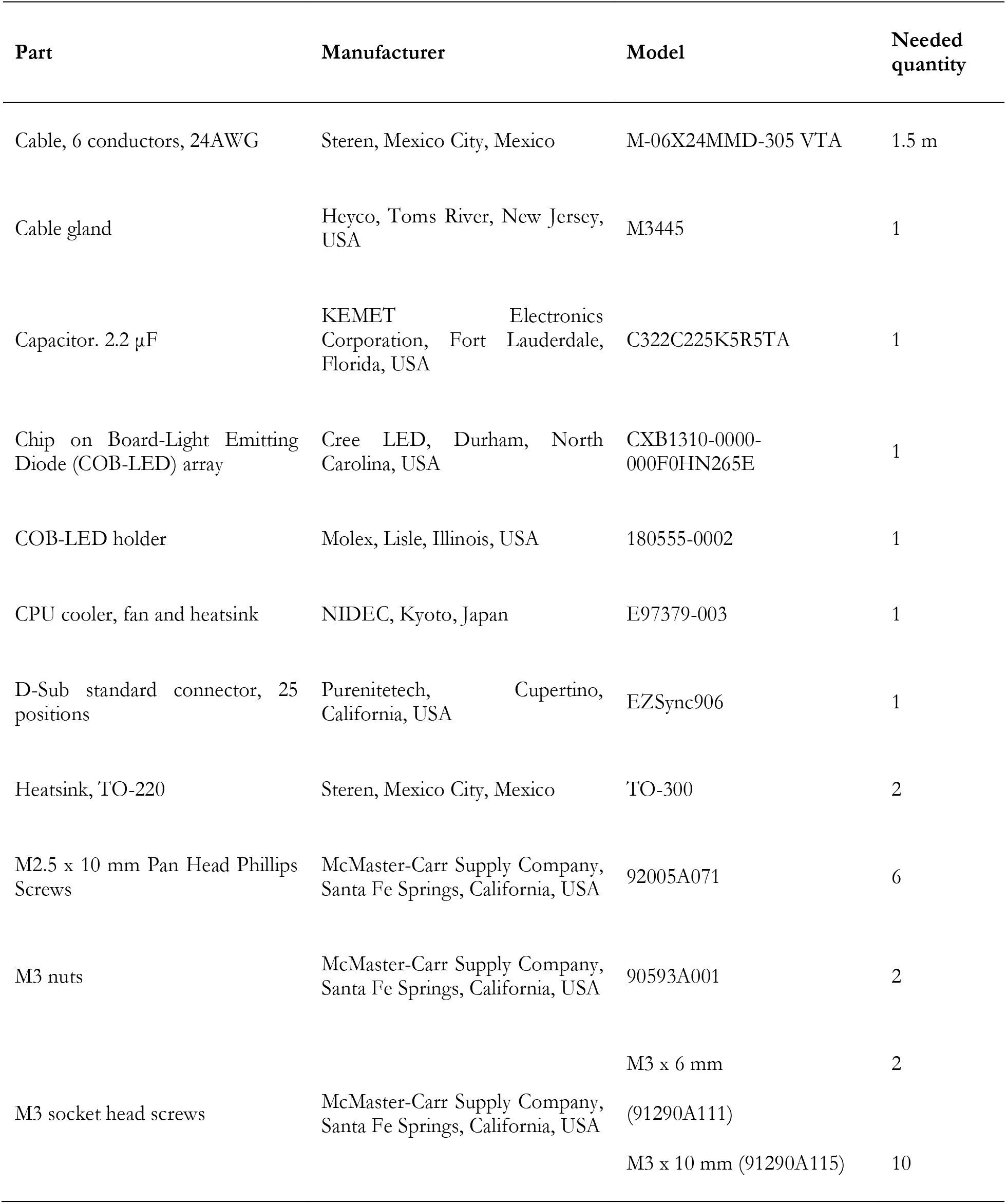
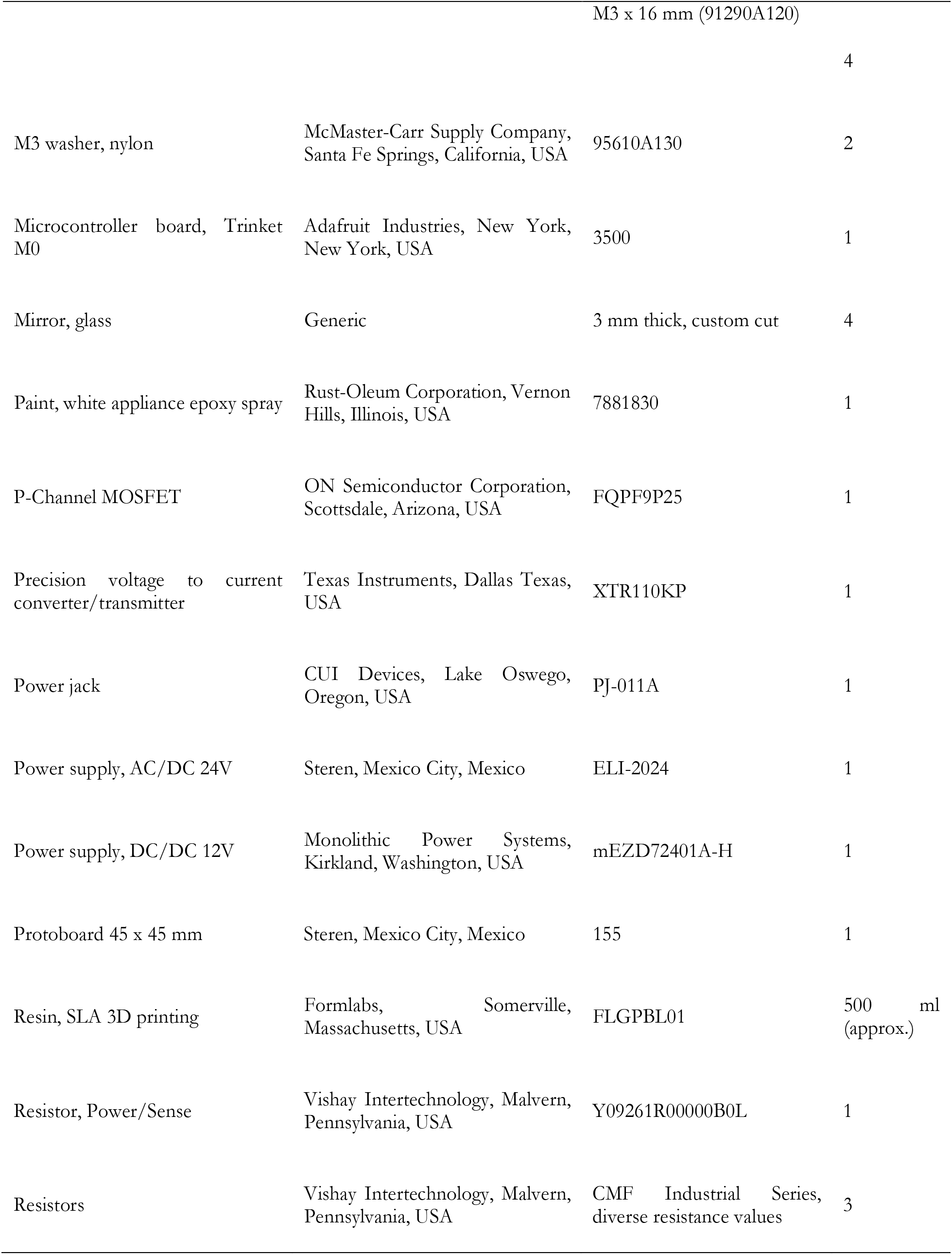
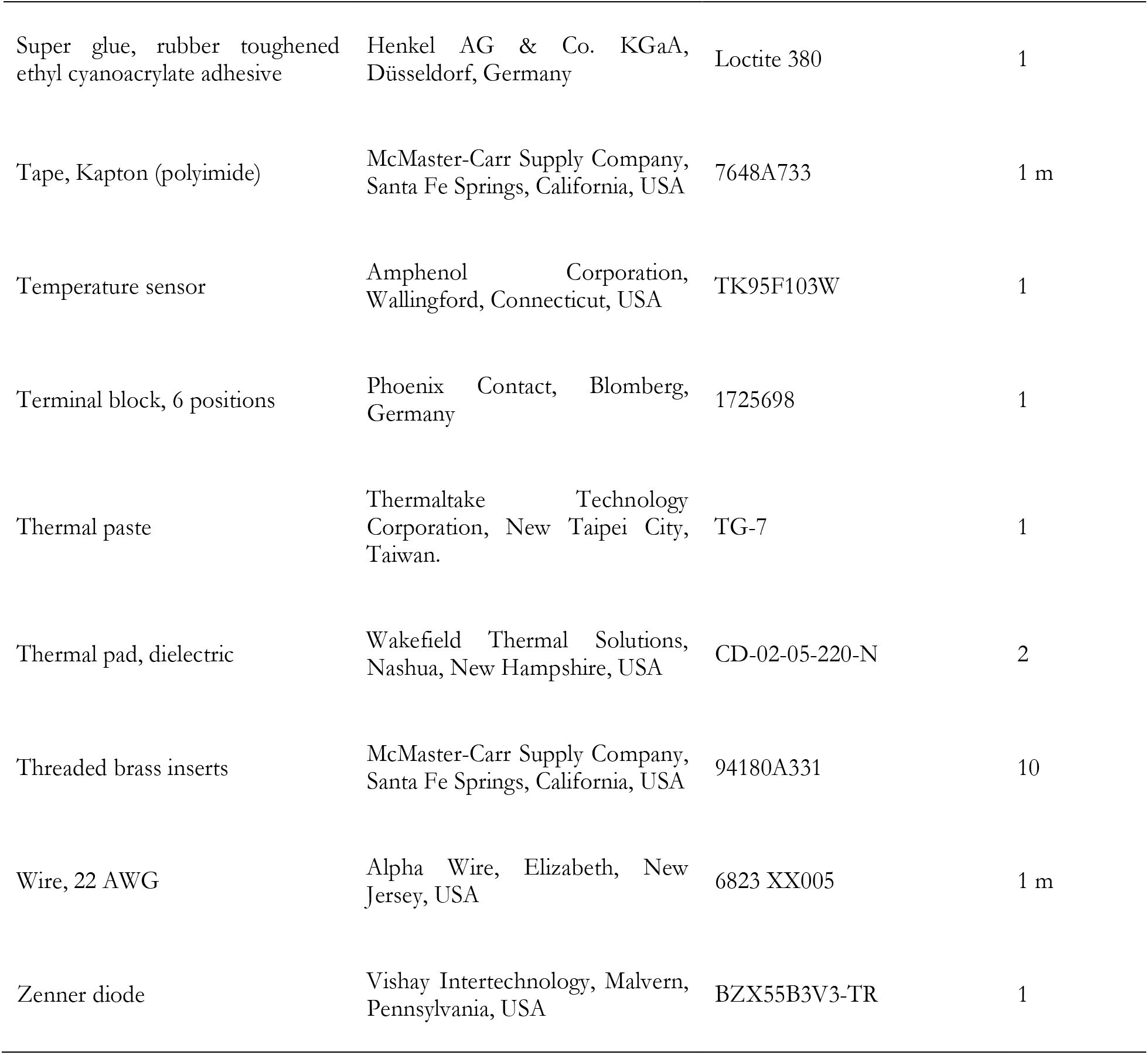
List of parts and materials needed to build the proposed LED lamp.

The current flows through the COB-LED array regulated by a custom-designed LED driver (Fig. 1). A detailed scheme of the electronic circuit is available on GitHub (https://github.com/AaronVelez/LI-6800_Lamp) under a CERN Open Hardware Licence Version 2, weakly reciprocal variant. The LED driver has three components. The first is a precision voltage-to-current converter/transmitter, the XTR110 chip (Table 1). The two other components, connected to the XTR110 chip, form a feedback circuit that monitors and controls the current to the COB-LED. A sense resistor monitors the current, and a P-channel metal oxide semiconductor field effect transistor (P-MOSFET) controls the current. Current from an external 24 V power supply flows through the sense resistor; a voltage drop across the resistor, proportional to the current, is measured by the XTR110 chip via the SOURCE-SENSE pin. In turn, the XTR110 chip adjusts the voltage to the P-MOSFET gate via the GATE_DRIVE_MOSFET pin. The sense resistor and the P-MOSFET transistor are both mounted on individual heatsinks. The regulated current flowing to the COB-LED array is proportional to the input voltage to the LED_CTRL pin; this control voltage is set by the LI-6800 console and has an operating range from 0 to 5 V.

**Figure 1.**
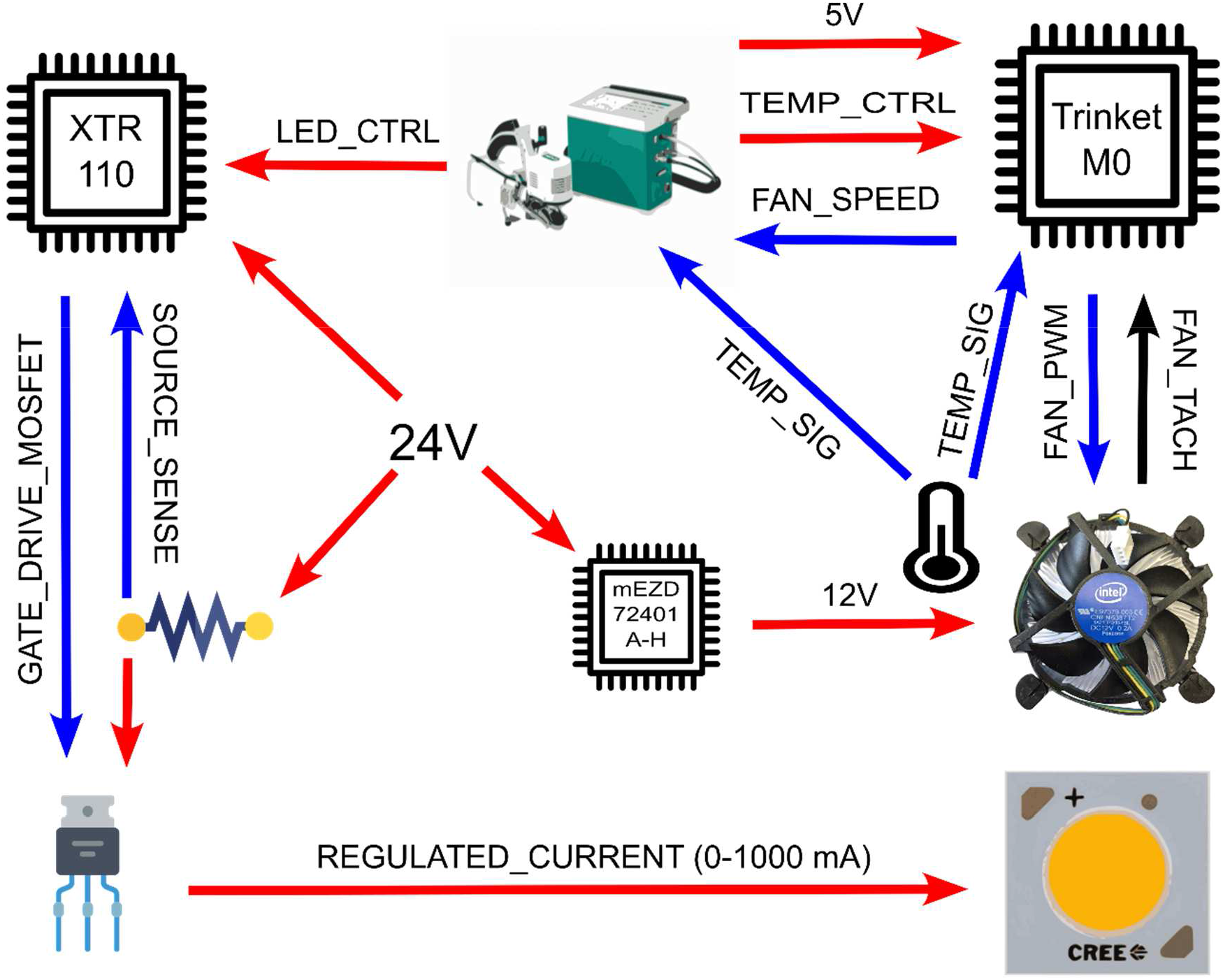
Simplified electronic design of the lamp for the LI-6800 photosynthesis system. Chip on Board-Light Emitting Diode (COB-LED) current is controlled by the XTR110 chip. Fan speed is controlled by the Trinket M0 microcontroller board in response to heatsink temperature. The COB-LED current and heatsink temperature setpoints are set by the LI-6800 via analog signals. Both the LI-6800 and the Trinket M0 monitor heatsink temperature using an negative temperature coefficient (NTC) thermistor. The Trinket M0 measures fan speed by monitoring the fan tachometer signal (FAN_TACH); then, the Trinket M0 translates the calculated fan speed into an analog signal and sends it to the LI-6800. An external 24 V AC-DC power supply powers an mEZD72401A-H DC-DC voltage converter, which in turns powers the XTR110 chip, COB-LED and fan with 12V. Red arrows indicate power sources; black arrows indicate digital signals, and blue arrows analog signals. The complete electronic schematics is available at the GitHub repository.

**Figure 2.**
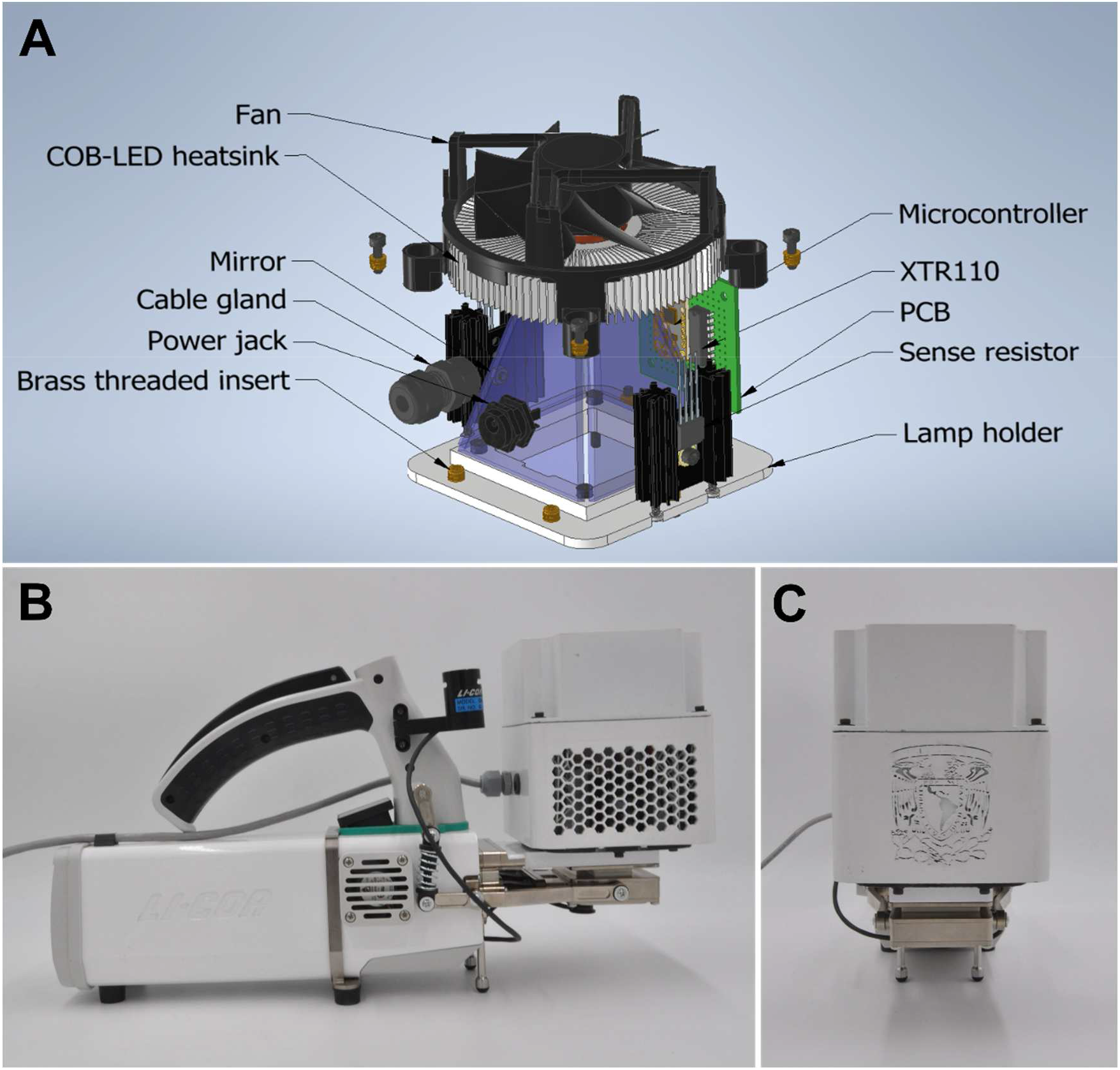
Mechanical design of lamp for the LI-6800 photosynthesis system. In panel (**A**), 3D assembly shows names of main internal parts. Notice that the main body and fan lid are not shown to depict internal parts. The complete 3D assembly is available at the GitHub repository. The lamp, attached to the LI 6800 sensor head is depicted in panels **B** (side view) and **C** (front view).

#### Temperature control and monitoring

To diminish temperature-dependent changes in PPFD output, the COB-LED array is mounted on a standard CPU cooler, which consists of a heatsink and a 4-wire fan (Table 1). A precision negative temperature coefficient (NTC) thermistor measures the internal heatsink temperature. The thermistor signal (TEMP_SIG) is measured by both the LI-6800 console and an Arduino-compatible microcontroller board. The microcontroller board we used is a Trinket M0 board manufactured by Adafruit (Table 1). The board main processor is an ATSAMD21E18 32-bit Cortex M0+ running at 48 MHz. It can be programed in C/C++ or Python. In this project, Python was used (see Software section). This board was selected because is only 27 x 15.3 mm in size. The microcontroller monitors the heatsink temperature and controls the fan speed.

The microcontroller uses the measured temperature to adjust the fan speed according to a proportional-integral-derivative (PID) algorithm that calculates the speed needed to reach the setpoint. This regulation occurs via a 25 kHz pulse-width modulation (PWM) signal (FAN_PWM). The microcontroller measures the actual fan speed from the tachometer digital signal on the FAN_TACH pin; then it encodes it into an analog signal and sends it to the LI-6800 console via the FAN_SPEED signal. The heatsink temperature setpoint is defined by an analog signal from the LI-6800 console (TEMP_CTRL). The LI-6800 can generate analog signals from −5 to 5 V; however, the microcontroller can only receive analog signals from 0 to 3.3 V. Protecting the microcontroller from potential damage by overvoltage, a Zener diode limits the TEMP_CTRL voltage to 3.3 V. The Zener diode is not pictured in Fig. 1 (see detailed electronic schematics at GitHub). The lamp design has no hardware or software protection from improperly setting TEMP_CTRL voltage to values lower than 0 V; if this is done, the microcontroller would be permanently damaged.

In addition to the microcontroller, the LI-6800 console measures the heatsink temperature by probing the same thermistor signal (TEMP_SIG). The LI6800 console uses the measured temperature to fine-adjust the voltage (LED_CTRL) needed to reach the light intensity required by the user. The effect of heatsink temperature on PPFD output was measured experimentally, and calibration coefficients were calculated using the collected data. See the following sections for details.

#### Electronic protoboard, interface and power supply

The XTR110 chip, microcontroller, a DC-DC converter, and diverse discrete electronic components were breadboarded on a 45 x 45 mm protoboard with standard 0.1 inches pitch (see Appendix S1). The sense resistor, P-MOSFET, and COB-LED generate heat that must be dissipated and were, therefore, placed on individual heatsinks. A shielded multiconductor cable with six individual conductor lines connects the lamp and the LI-6800 console. At the LI-6800 console side, a 25 position D-Sub standard connector attaches the individual signal lines to the *User I/O* port. At the lamp side, a PCB terminal block attaches the induvial conductor lines to the protoboard.

The lamp is powered by two independent sources. First, the COB-LED driver and cooling fan are powered from an external 48 W, 24 V AC-DC power supply. The COB-LED driver is designed to use 24 V, but the fan needs 12 V. Therefore, a DC-DC converter reduces the voltage from 24 to 12 V and delivers power to the fan. Without considering inefficiencies, the COB-LED driver and fan, at maximum power, consume at least 20.4 W. Second, the microcontroller is powered from the LI-6800 console via the *User I/O* pin 14 (*Power5*). This line delivers 5 V when it is turned ON from the LI-6800 console.

#### Lamp housing and assembly

A 3D-printable housing was initially designed with SolidWorks (Dassault Systèmes SE, 2016) and later refined with Autodesk Inventor (Autodesk Inc., 2019). The lamp housing is composed of three parts, the main body, fan lid and lamp holder. The main body holds two heatsinks (carrying the P-MOFTET and the sense resistor), the electronic protoboard (carrying the microcontroller and LED driver), the CPU cooler assembly (carrying the COB-LED and the thermistor), and four mirrors. The fan lid prevents the user from touching the fan blades and holds the CPU cooler assembly in place. The lamp holder fixes the lamp housing to the LI-6800 sensor head. The lamp holder is compatible with the Clear-top Leaf Chamber, model 6800-12A from Li-Cor Biosciences. Editable 3D models, and ready-to-print files are available at GitHub.

The lamp housing parts were printed with a stereolithography (SLA) 3D printer (model From 2) using Black resin v1.0 (Formlabs, Somerville, Massachusetts, USA). After 3D printing, parts were first cleaned with isopropyl alcohol using the From Wash accessory equipment (Fromlabs); then, parts were cured at 60 °C for 30 min using the From Cure accessory equipment (Fromlabs). Later, the parts were painted with epoxy aerosol paint. Finally, heat-set threaded brass inserts were fixed at the top and bottom of the main body to fix the fan lid and lamp holder using M3 screws. Table 1 shows the model and manufacturer of all used parts. As of September 2023, the approximate cost for all materials and parts is 260.95 USD. Appendix 1 gives step-by-step fabrication instructions.

### Lamp and LI-6800 software

#### Microcontroller code

The used microcontroller is pre-loaded with CircuitPython (Adafruit Industries, 2023), which is a beginner friendly, open-source version of Python 3. In turn, CircuitPython is a derivative from MicroPython (The MicroPython project, 2023). The microcontroller monitors and controls the COB-LED heatsink and its fan using standard libraries, including *board*, *pulseio*, *time*, *math*, *digitalio* and *analogio*. As CircuitPython does not have native PID algorithms, we programed a simple PID controller using only basic functions. Our PID controller is based on published code (Kantor, 2020). The microcontroller python script used in the lamp is available at GitHub, and it is published under a GNU Lesser General Public License v3.0.

The code loaded to the microcontroller is a simple loop with five steps. The first step reads the analog signal at the TEMP_CTRL pin, which encodes the COB-LED heatsink temperature setpoint defined by the LI-6800 console. Step two measures the actual COB-LED temperature. This is done by measuring the voltage at the TEMP_SIG pin and calculating the thermistor resistance using Ohms Law and the known thermistor excitation voltage (3.3V). Then, thermistor resistance is translated to thermistor temperature using manufacturers temperature-resistance equations and coefficients. At step three, the fan speed is read at the FAN_TACH pin. Then, the microcontroller encodes the fan speed into an analog signal and sends it to the LI-6800 console at the FAN_SPEED pin. During step four, the PID algorithm is computed; it calculates the PWM duty cycle needed to reach the COB-LED heatsink temperature setpoint. Finally, during step five, the calculated PWM duty cycle is outputted to the FAN_PWM pin.

#### LI-6800 custom configuration and background program

The lamp COB-LED driver and microcontroller get the PPFD and temperature setpoints from the LI-6800 console *User I/O* channels DAC1 and DAC2, respectively. That is, DAC1 is connected to the LED_CTRL pin and DAC2 to the TEMP_CTRL pin (see *User I/O* D-Sub connector pin mapping at the LI-6800 manual). Although the lamp can be controlled by changing DAC1 and DAC2 voltage (Fig. 3A), adjusting the values manually would be impractical. This is because the mapping between DAC voltage and PPFD or temperature setpoints depends on calibration coefficients. To ease the lamp control by the user, therefore, we developed a custom LI-6800 configuration file and programed an LI-6800 BP named *PAR_Ctrl.py*. The custom configuration file and BP can be downloaded from GitHub.

**Figure 3.**
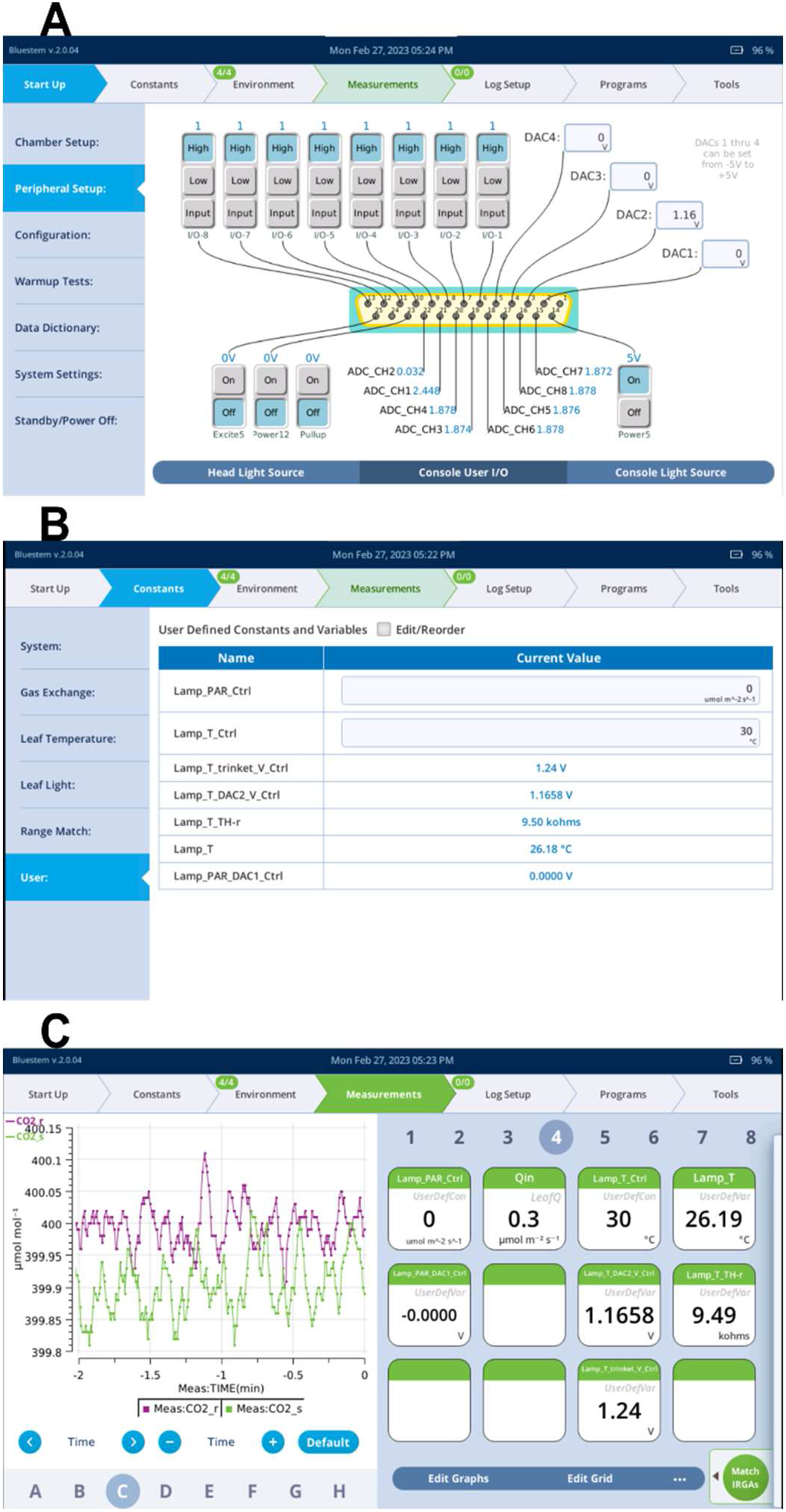
LI-6800 screens while using the lamp. Panel (**A**) shows the custom *User I/O* state loaded from the configuration file. Notice that the 5 V power must be ON in order to power the microcontroller. Panel (**B**) shows the user constants and variables added by the configuration file. Notice that the user constants Lamp_PAR_Ctrl and Lamp_T_Ctrl are settable fields to ease lamp use. The non-settable fields are user variables and make all necessary calculations to control the lamp programmatically. Panel (**C**) shows that the configuration file also adds user constants and variables to the grid display in the *Measurements* tab.

The custom LI-6800 configuration file has only three configuration fields. Those are user definitions (*UsrDef*), the configuration of variables at the *Measurements* tab (*Grids*), and the state of the *User I/O* channels (*Aux*). Notice that *User I/O* was formerly known as *Auxiliary I/O*; in the LI-6800 Bluestem OS v2.0 manual, the term *Auxiliary I/O* is still used. In this study, we used the Bluestem OS v2.0.04 that holds the new term *User I/O*.

When the *UsrDef* field is loaded from the configuration file, user constants and variables (*UsrDef*) are added. User-defined constants are shown at the LI-6800 Console *Constant > User* tab as user-settable fields (Fig. 3B).The user, therefore, can input the desired COB-LED PPFD and temperature setpoints in the target units; those are μmol·m^−2^·s^−1^ and °C for the PPFD and temperature, respectively. In turn, the loaded user variables calculate the DAC1 and DAC2 voltages that are required to reach the desired setpoints. The equations and calibration coefficients needed to calculate these conversions are within the configuration file (*UsrDef* field). Notice that the user variables only calculate the required DAC1 and DAC2 voltages, but they do not set those voltage values in LI-6800 *Console User I/O* tab. Nonetheless, the *PAR_Ctrl.py* BP automatically sets DAC1 voltage to the required voltage. The *PAR_Ctrl.py* BP needs to be run by a user or by another BP. Once started, *PAR_Ctrl.py* BP runs in the background, allowing the user to control the lamp with ease. As the user variables are also settable by any BP, the lamp can be fully controlled programmatically.

When the *Aux* field is loaded from the configuration file, the 5 V power supply from the *User I/O* is turned ON (pin 14 in the D-Sub connector) and powers the microcontroller. The *Aux* field also sets the DAC2 voltage to 1.1658 V, which corresponds to a COB-LED heatsink temperature setpoint of 30 °C. This prevents the user forgetting to turn ON the microcontroller and setting the heatsink temperature. The LI-6800 *User I/O* channel ADC_CH1 measures the voltage at the TEMP_SIG pin. Using the read voltage and calibration coefficients, a user variable calculates the COB-LED heatsink temperature. Similarly, the ADC_CH2 channel measures voltage at the TEMP_SIG pin, which is used to calculate fan speed. When the *Grid* field is loaded from the configuration file, the variables associated with lamp operation are displayed at the LI-6800 console *Measurements* tab (Fig. 3C) for convenient monitoring during measurements.

### Lamp calibration and characterization

#### Lamp emitted light characterization and calibration

We measured the spectral distribution of the emitted light using a ultraviolet–visible, handheld, portable spectrophotometer with one nm spectral resolution over the range 400 to 1000 nm (Stellarnet, Tampa Florida, USA). The spectrophotometer was calibrated with a mercury-argon wavelength calibration source (Ocean Optics, Rochester New York, USA). The measurement was taken at the bottom of the lamp housing, directly below the COB-LED; that is, the spectrophotometer probe was at the same distance from the COB-LED as a leaf is during measurements. This ensures that any potential effects of the housing and mirrors on the spectral distribution are measured. The spectral measurement was done setting the lamp at 1% of its power and using an integration time of 10 ms. The measured spectrum is shown in Fig. 4A. The light emitted at the photosynthetically active range (between 400 and 700 nm) corresponds to 99.5% of the total emitted light. The COB-LED emits light with a red to far-red ratio of 11.99, and the calculated phytochrome photostationary state (PSS) (C. Sager et al., 1988) is 0.86.

**Figure 4.**
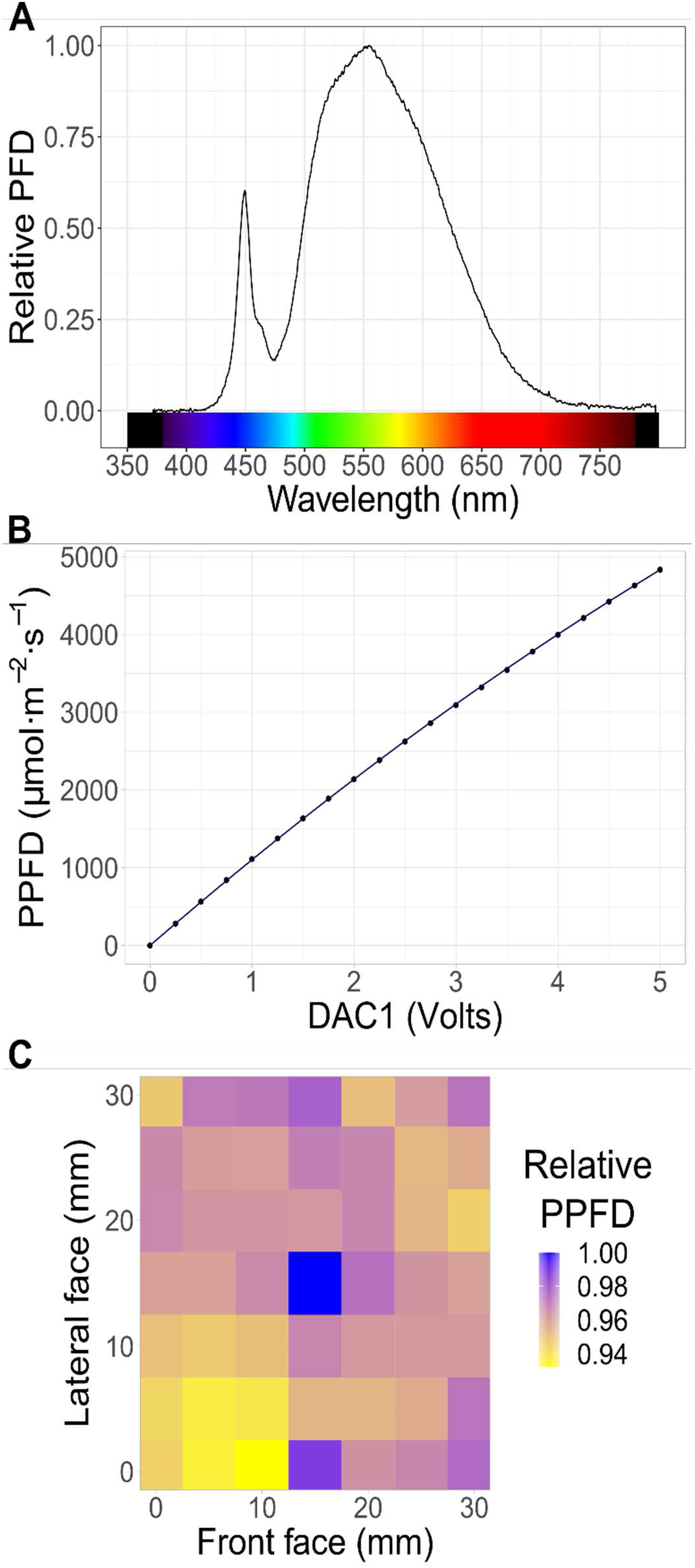
Characterization of the lamp for the LI-6800 photosynthesis system. Panel (**A**) shows the spectral distribution of light at the leaf plane. Panel (**B**) shows the resulting photosynthetically active photon flux density (PPFD) at the leaf plane as a result of DAC1 voltage. Panel (**C**) shows the relative PPFD distribution at the leaf plane using the 3 x 3 cm aperture insert.

The custom driver circuit increases the current to the COB-LED in response to the voltage at the LED-CRTL pin, which is connected to the LI-6800 DAC1 *User I/O* channel. In turn, the light emitted by the COB-LED is proportional to the current; however, as with any LED, there are also small effects of the temperature on the emitted light intensity. As the COB-LED temperature increases, the emitted light, at any given current, slightly decreases. The effect of the leaf chamber aperture inserts needs also to be considered. The leaf chamber (model 6800-12A) has three aperture inserts (3×3, 2×3 and 1×3 cm). When using an insert with a smaller aperture, more metal surface reflects the emitted light back to the mirrors — and then back to the leaf —, resulting in a slight increase in PPFD at the leaf surface. To accurately control the COB-LED, therefore, we performed a calibration that considers these three variables defining PPFD at the leaf surface. The detailed calibration procedure is outlined in Appendix 2. In summary, we recorded the COB-LED heatsink temperature and the actual PPFD inside the leaf chamber using a light meter (model LI-250A) equipped with a quantum sensor (LI-190R Quantum) (LI-COR Biosciences, Lincoln, Nebraska, USA) at DAC1 voltages from 0 to 5 V. This procedure was repeated two times with each of the three aperture inserts. As expected, PPFD increases in response to DAC1 voltage (Fig. 4B). Using SAS Studio (SAS Institute Inc., 2022), we evaluated several multiple linear regression models using DAC1, COB-LED heatsink temperature and aperture insert area as predictor variables. The selected model is as follows:

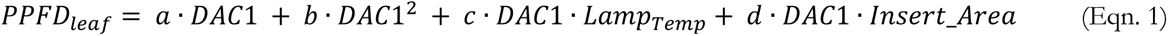

Where *PPFD_leaf_* is the predicted PPFD at the leaf surface within the leaf chamber in µmol·m^−2^·s^−1^. *DAC1* is the voltage at the LI-6800 *User I/O* DAC1 channel in volts. *Lamp_Temp_* is the COB-LED heatsink temperature in °C. *Insert_Area* is the aperture insert area in cm^2^; the 3×3 aperture inserts results in an *Insert_Area* 9 cm^2^. Letters *a*, *b*, *c* and *d* represent regression coefficients. Table 2 shows the estimated coefficients. The resulting r^2^ for the selected model and the used calibration data is 1.00. This means that the calibration equation can accurately predict the resulting PPFD at leaf surface. As the calibration equation and coefficients are included in user variables within a LI-6800 custom configuration file, the user can control the PPFD of any experiment by running the *PAR_Ctrl.py* BP and adjusting, manually or programmatically, a user constant.

**Table 2.**
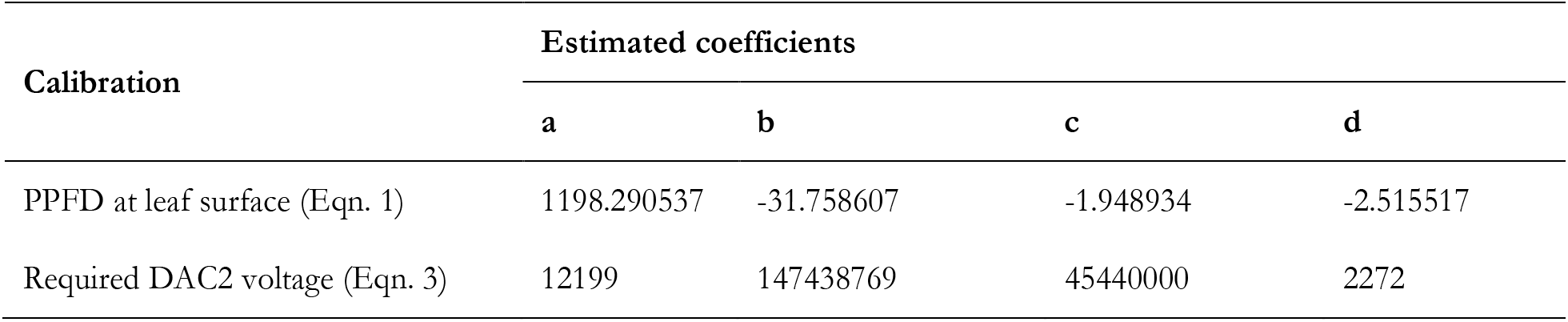
Calibration coefficients.

| Calibration | Estimated coefficients |  |  |  |
| --- | --- | --- | --- | --- |
|  | a | b | c | d |
| PPFD at leaf surface (Eqn. 1) | 1198.290537 | -31.758607 | -1.948934 | -2.515517 |
| Required DAC2 voltage (Eqn. 3) | 12199 | 147438769 | 45440000 | 2272 |

In addition to the calibration defining the relation between DAC1 control voltage and PPFD at leaf level, we developed a way the user can measure PPDF during an experiment by using the photodiode sensor built into the leaf chamber. According to the LI-6800 manual, the built-in photodiode sensor reading is used to calculate the variable *Q_amb_in_* using the following equation:

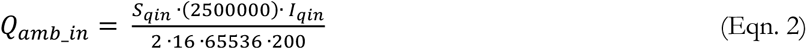

Where *Q_amb_in_* is the ambient light PPFD inside the leaf chamber in µmol·m^−2^·s^−1^. *I_qin_* is the photodiode raw counts from the ADC converter. The calibration factor *S_qin_* is stored within the leaf chamber memory as a multiplier with units µmol·m^−2^·s^−1^·µA^−1^. Without re-calibration of the *S_qin_* multiplier, *Q_amb_in_* overestimates the COB-LED PPFD output (see Appendix S2). This is probably because the built-in photodiode is placed closer to the COB-LED than the external quantum sensor, which was placed at the leaf plane. Up to a PPFD of around 2700 µmol·m^−2^·s^−1^, however, there is a perfect linear correlation between *Q_amb_in_* and the measured PPFD using an external quantum sensor, resulting in an r^2^ = 1.00. Above 2700 µmol·m^−2^·s^−1^, *Q_amb_in_* signal saturates. This means that the built-in photodiode can be used to monitor PPFD up to 2700 µmol·m^−2^·s^−1^ if recalibrated with a new *S_qin_* value. The new *S_qin_* value was calculated by dividing the original *S_qin_* by the slope of the linear relation between *Q_amb_in_* and the measured PPFD using the external quantum sensor. The estimated *S_qin_* is 6.40736 µmol·m^−2^·s^−1^·µA^−1^. The new multiplier results in a perfect linear relation between *Q_amb_in_* and the PPFD value measured with an external quantum sensor (Appendix S2).

Once the built-in photodiode is recalibrated, *Q_amb_in_* could be used to feedback PPFD control by Eqn. 1. Although, after a calibration, Eqn. 1 is enough to control PPFD, further DAC1 voltage fine adjustment by *Q_amb_in_* feedback can compensate for the decrease in COB-LED intensity associated with hours of use. However, such a decrease is slow. According to manufacturer tests, the used COB-LED retains 90% of its irradiance output even after 66500 h of use at maximum power (Cree LED, 2023). Additionally, not all leaf chambers are equipped with an internal photodiode. Therefore, we developed two ways to control PPFD, which are with and without *Q_amb_in_* feedback. Both control approaches have advantages and disadvantages. The approach not using feedback allows the lamp to be used on any chamber and at any intensity, but it requires recalibration as the COB-LED gets older. Using the feedback approach requires less frequent recalibration as the COB-LED gets old; however, it will not work with all leaf chambers, and it only would work at a PPFD lower than 2700 µmol·m^−2^·s^−1^. We developed scripts and step-by-step instructions to use either approach (Appendix 4).

As the lamp housing holds four mirrors and places the COB-LED 68.7 mm above the leaf chamber top surface, the emitted light is expected to homogenously illuminate the leaf. To corroborate this prediction, we measured the light distribution at the leaf plane. Using the external quantum sensor and a XY platform (Model Zyshtnc XY125C, Dongguan Fara Precision Parts Co., Jiangsu, China) we measured PPFD at 49 locations (7 x 7 XY grid) while setting DAC1 to 1 V. The resulting light distribution is shown in Fig. 4C; the reported values are relative to the PPFD at the center of the leaf chamber, which was 1484.3 µmol·m^−2^·s^−1^. The maximum spatial variation in PPFD was 6.7%, and most grid quadrants (81.63%) had a variation lower than 5%.

#### Lamp temperature control characterization and calibration

The COB-LED heatsink temperature setpoint is set by the voltage at the TEMP_CTRL pin, which is connected to the LI-6800 DAC2 *User I/O* channel. As stated in the hardware section, however, a Zener diode reduces the voltage generated by the LI-6800 to protect the microcontroller circuits. Using a multimeter (model 87V, Fluke Corporation, Everett, Washington, USA), the voltage reduction by the Zener diode was measured (See Appendix S3). To define the DAC2 voltage required to inform the microcontroller the temperature setpoint, these measurements were fitted to the following equation using the *curve_fit* function of the *scipy* Python library.

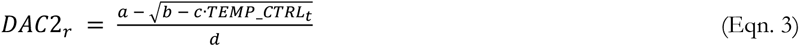

Where *DAC2_r_* is the DAC2 voltage required to reach the target voltage at the microcontroller side (*TEMP_CTRL_t_*). Letters *a*, *b*, *c* and *d* represent the regression coefficients. Table 2 shows the estimated coefficients. In turn, *TEMP_CTRL_t_* is defined by the following equation:

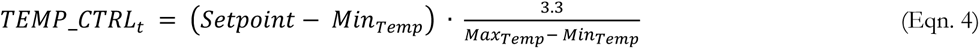

Where *Setpoint* is the user-defined COB-LED heatsink temperature setpoint. The variables *Min_Temp_* and *Max_Temp_* are the minimum and maximum, selectable temperature setpoints, respectively. Equations. 3 and 4 are programed within the LI-6800 custom configuration file, while the microcontroller software only needs Eqn. 4. Within the microcontroller software and custom LI-6800 configuration file, *Min_Temp_* and *Max_Temp_* are set at 15 and 55 °C, respectively.

Although the Lamp software allows the user to set a heatsink temperature setpoint between 15 and 55 °C, those temperatures are not always reachable. To characterize the ability of the fan and heatsink to reach the setpoint temperature, we varied the COB-LED intensity from 0 to 100% and changed the heatsink temperature setpoint from 2 °C lower up to 14 °C higher than the room temperature. However, we did not try all combinations of COB-LED intensity and temperature setpoint. For example, at high COB-LED intensity, it is not feasible to use a setpoint close to room temperature. Hence, we selected 884 combinations that were closest to the attainable range. The results are shown in Appendix S4. At maximum COB-LED intensities, the fan does not cool the heatsink to a temperature less than 10 °C above room temperature; however, even if the heatsink heats up to 80 °C, the attainable PPFD is 4890 µmol·m^−2^·s^−1^ (Appendix S5). This PPFD is more than enough to perform photoinhibition experiments, usually done between 3000 and 3700 µmol·m^−2^·s^−1^, for example (Velez-Ramirez et al., 2017). Nonetheless, most plant sciences experiments would require a PPFD of less than 2000 µmol·m^−2^·s^−1^, for example, an A-PPFD curve (Coe and Lin, 2018). A PPFD of 2000 µmol·m^−2^·s^−1^ is reached at only 40% of the COB-LED intensity (Fig. 4B and Appendix S5). Consequently, at a COB-LED intensity up to 40%, a heatsink temperature setpoint up to 2 °C higher than room temperature is adequate for most cases (Appendix S4).

### Lamp example usage

To demonstrate the use of the lamp, we performed A-PPFD curves using wheat (*Triticum aestivum*) plants. Wheat seeds, Borlaug100 F2014 variety, were kindly provided by the International Center for the Improvement of Maize and Wheat (CIMMYT, Texcoco, Mexico). Seeds were sown directly on fresh soil mix on March 4^th^. Each batch of soil mix contained 250 L of peatmoss (REMIX 1) (Rėkyva Joint-Stock Company, Šiauliai, Lithuania), 200 L of perlite (Multiperl Hortícola) (Grupo Perlita, Gómez Palacio, Durango, Mexico), 100 L of vermiculite (Disa Sustratos Agrícolas, Mexico City, Mexico), 100 L of coconut fiber (Hydro Environment, Tlalnepantla, Mexico), 60 L of silica sand (Decoproductos, Mexico City, Mexico), 2 kg of dolomite limestone (Zeolitech, Cuernavaca, Morelos, Mexico), 2 kg of slow-release N-P-K (16-16-16) fertilizer (Yara Mila UNIK16) (Yara, Oslo, Norway), 875 g of chelated micronutrient mix (Kelatex Multi) (Cosmocel, Monterrey, Nuevo León México), 375 g of magnesium sulphate heptahydrate (Industrias Peñoles, Mexico City, Mexico), and 150 g of calcium sulphate (Grow Depot Mexico, Torreón, Coahuila, Mexico). Plants were grown in a glass greenhouse located within the University facilities in the city of León, Guanajuato, México. When plants were 24 days old, the youngest, fully expended leaf was measured.

Using the lamp presented here, A-PPFD curves were done with LI-6800 photosynthesis measuring system (Li-Cor Biosciences). A python script was used to perform the measurements automatically. The script can be found at the GitHub repository, and Appendix 4 details instruction on how use it. Next, a brief description of the environmental conditions is provided. Leaf temperature was controlled at 25 °C. Flow rate was set at 275 µmol·s^−1^ when the PPFD was lower than 400 µmol·m^−2^·s^−1^ and to 350 µmol·s^−1^ when PPFD was higher than 400 µmol·m^−2^·s^−1^. Fan speed was set at 9000 rpm. Carbon dioxide concentration at the reference infrared gas analyzer (IRGA) was set at 400 ppm, and the vapor pressure deficit (VPD) at 1.2 kPa. Assimilation rate was measured at ten PPFD levels, which were 0, 41, 93, 147, 332, 516, 900, 1274, 1878 and 2459 µmol·m^−2^·s^−1^. After each change in PPFD, the script waited for stability before logging. A match of the reference and sample IRGAs was done before each log. Figure 5 shows the resulting A-PPFD curves. Photosynthetic parameters were estimated using a rectangular hyperbola model (Rivera-Méndez and Romero, 2017) according to the following equation:

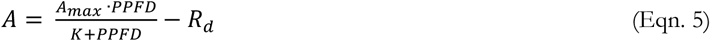

**Figure 5.**
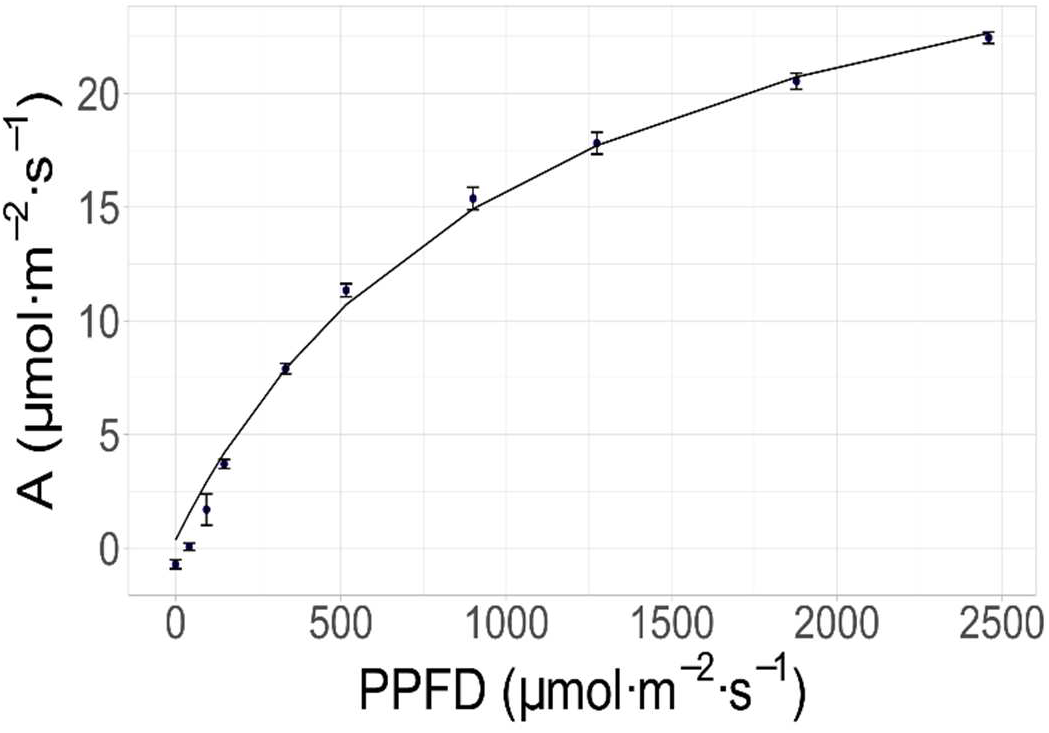
Lamp example usage. Using the lamp developed here, an A-PPFD curve was measured. Each point is the average of four biological replicates. Error bars show the standard error of the mean. Using the model depicted in Eqn. 5, the regression line is the result of fitting all measurements in a single fit procedure. The calculated photosynthetic parameters are reported in Table 3.

**Table 3.** Photosynthetic parameters.

| Parameter | mean | SE |
| --- | --- | --- |
| $R_d$ | -1.82 | $\pm 0.14$ |
| $A_{max}$ | 30.99 | $\pm 0.24$ |
| $K$ | 506.23 | $\pm 22.92$ |

Where *A* and *A_max_* are the CO_2_ assimilation rate and maximum assimilation rate, respectively. *PPFD* is the incident irradiance. The parameter *K* is the light saturation constant, and *R_d_* is the dark respiration rate. All units are in µmol·m^−2^·s^−1^. All the parameters were estimated using Microsoft Excel. Using only the measurements at limiting light, *R_d_* was estimated as the slope of a linear regression between A and PPDF. The parameters *A_max_* and *K* were estimated by minimizing the least square error. Table 3 shows the photosynthetic parameter estimates.

## CONCLUSIONS

Here, we report the design, construction, testing and use of a lamp compatible with the LI-6800 photosynthesis system. The lamp can be built using a basic electronics lab and a 3D printer. We share calibration instructions, which only require equipment commonly available at plant science laboratories. The lamp is a cost-effective alternative to perform photosynthesis research coupled with the popular LI-6800 photosynthesis measuring system.

## AUTHOR CONTRIBUTIONS

AIVR, JdDM and HCA took part in the hardware design. AIVR, JdDM and UGPG programed the required software. AIVR, JdDM, UGPG and AMJ performed the lamp characterization and calibration. AIVR, JdDM and UGPG participated in the biological measurements. AIVR, UGPG and JVM supervised BSc. students working on the project. AIVR, JVM and IHM conceptualized the idea. AIVR, JVM and IHM drafted the paper. AIVR and IHM secured the funding. All authors reviewed and approved the manuscript.

## ACKNOWLEDGMENTS

The authors thank for the funding to CONACYT Infrastructure project 294511, and UNAM-PAPIIT projects IA207519 and IA207021. We also thank to UNAM-PAPIIT for the BSc thesis scholarship to JdDMG and CONACYT for the full MSc scholarship to UGPG.

## DATA AVAILABILITY

The data and metadata from the A-PPFD measurements are available at the GitHub repository. The reporting format follows the recommendations defined for leaf-level gas exchange data (Ely et al., 2021).

## Appendix 1. Lamp construction

This section provides step-by-step instructions on how to build a lamp for the LI-6800 photosynthesis measuring system. Table 1 details every part and material required. The CAD files and electronic schematics are available at GitHub (https://github.com/AaronVelez/LI-6800_Lamp). This section provides printing instructions using the STL files; however, the original files in IPT and STP formats are also provided in case the user needs to modify the CAD design. The provided instructions apply for the Form 2 3D printer from FromLabs; however, the protocol can be adjusted for other 3D printers.

### 3D Printing of the Lamp housing

1. From the GitHub repository, download the *Main Body.stl*, *Fan Lid.stl* and *Lamp Holder.stl* files.
2. Download and install the PreForm software https://formlabs.com/software/
3. Open the *Main Body.stl* file with PreForm.
4. Orient the part so its base sits on top of the printing platform.
5. Generate 3D printing supports using default settings.
6. At the *Job Setup* menu, select printer (Form 2), material (Black resin) and the z axis resolution (50 µm).
7. Print the job.
8. Remove the printing platform from the printer and wash the resin residues with isopropyl alcohol. If available, use the *From Wash* equipment (FormLabs); otherwise, do it manually.
9. Remove the part from the printing platform. Do not remove the 3D printing supports yet.
10. Cure the part using the *Form Cure* equipment (FormLabs). Set the curing temperature and time to 60 °C and 30 min, respectively. Make sure to place the part inside the equipment before it warms up to prevent deformations.
11. Remove the 3D printing supports using pliers.
12. Using wet sandpaper, smooth the part surfaces. Start with coarse sandpaper and gradually use smoother sandpaper; for example, use sandpaper with grit numbers 120, 320 and 600, in that order.
13. Paint the part exterior with epoxy paint. Do not paint the interior. Apply fine layers, allowing each layer to dry before applying the next one. Three layers are recommended.
14. Repeat steps 3 to 13 with the *Fan Lid.stl* part.
15. Repeat steps 3 to 12 with the *Lamp Holder.stl* part; that is, do not paint this part as it would prevent proper fit within the *Main body* part.

### Adding Threads

The following instructions assume that you use the exact threaded brass inserts and heatsinks as indicated in Table 1. Adjust drilling and tapping bits if alternative threaded inserts and/or heatsinks are used.

1. From the GitHub repository, download the *Lamp.iam* assembly file as well as all individual parts files (all *\*.ipt* files).
2. Visualize the Lamp assembly using AutoDesk Viewer online tool at https://viewer.autodesk.com/designviews.
3. By exploring the assembly with the online viewer, identify the places of the Lamp main body where brass threaded inserts need to be installed. The spots are marked with 3 mm holes, and there are four places on top and six at the bottom.
4. At the identified spots, drill a 4 mm depth hole using a number 8 drill bit (5.0546 mm in diameter).
5. Install the threaded inserts by pressing them into place with a hot soldering tip. If brass inserts fall after installation, use a drop of superglue to secure them again to the lamp main body.
6. By exploring the assembly with the online viewer, identify the places of the TO-220 heatsinks where threads need to be tapped.
7. At the identified spots, machine an internal thread using a M2.5 tapping bit.
8. By exploring the LED holder datasheet (https://tools.molex.com/pdm_docs/sd/1805550002_sd.pdf), identify the places of the CPU cooler heatsink where threads need to be tapped.
9. At the identified spots, drill with the adequate drill bit before machining an internal thread using a M2.5 tapping bit.

### Protoboarding electronic components

1. From the GitHub repository, download the *Lamp_schematics.pdf* file, which contains detailed electronic schematics.
2. Download the Supporting information, containing the Appendix S1.
3. Using images in Appendix S1 as a guide and following the schematic design, protoboard all electronic components in a PCB. Notice that the P-channel MOSFET and the Power/Sense resistor are mounted on the heatsinks rather than on the PCB.
4. Using 22 AWG wire, solder the P-channel MOSFET and the Power/Sense resistor to the PCB.
5. Using the included lead wires, solder the fan assembly and the COB-LED holder to the PCB.

### Lamp assembly

1. From the GitHub repository, download the *Lamp.iam* assembly file as well as all individual parts files (all *\*.ipt* files).
2. Visualize the lamp assembly using AutoDesk Viewer online tool at https://viewer.autodesk.com/designviews
3. By exploring the assembly with the online viewer, identify the placement of all components.
4. Solder 22AWG wire leads to the power jack.
5. Install the power jack and cable gland in the lamp main body.
6. Solder the power leads to the PCB.
7. Install the P-channel MOSFET and the Power/Sense resistor onto the TO-220 heatsinks, using M3 x 6 mm screws. Make sure you use dielectric thermal pads and nylon washers to electrically isolate the heatsink from the electronic components.
8. Secure the TO-220 heatsinks to the lamp main body using M2.5 screws.
9. Insert the multiconductor cable thought the cable gland.
10. Connect the individual conductors to the PCB terminal block.
11. Connect the individual conductors on the other end of the cable to the D-Sub standard connector.
12. Place the PCB into the slots inside the lamp main body.
13. Remove the fan from the CPU cooler heatsink.
14. Remove the clips to attach the fan frame to a computer motherboard.
15. Using a 3 mm drill bit, drill a hole through the CPU cooler heatsink from top to bottom.
16. Place thermal paste onto the CPU cooler heatsink.
17. Secure the COB-LED to the CPU cooler using the COB-LED holder and 2.5 screws.
18. From the top of the CPU cooler heatsink, fill the hole drilled in step 14 with thermal paste.
19. Insert the temperature probe (NTC thermistor) in the hole filled with thermal paste so it touches the back plate of the COB-LED array.
20. Use a piece of Kapton tape to secure the temperature sensor wires to the heatsink.
21. Place back the fan on top of the CPU cooler heatsink.
22. Make sure all wires are nicely arranged away from the COB-LED.
23. Tighten the cable gland.
24. Place the CPU cooler on top of the lamp main body.
25. Place the fan lid on top of the fan and secure it with M3 x 10 mm screws.
26. Insert the mirrors in their designated slots in the underpart of the lamp main body. It is recommended to secure the mirrors with a piece of folded paper inserted between the mirror and the lam main body.
27. Using M3 x 16 mm screws, attach the lamp holder to the LI-6800-12A Clear-top Leaf Chamber.
28. Place the lamp assembly on top of the lamp holder and secure it using M3 x 10 mm screws.
29. Connect the D-Sub standard connector to the LI-6800 *User I/O* port.
30. Connect the AC-DC power supply to the lamp power jack.

### Microcontroller software installation

1. From the GitHub repository, download the *Microcontroller_Main.py* file.
2. Install the latest CircuitPython version in the Trinket M0 microcontroller following manufacturer instructions at https://learn.adafruit.com/adafruit-trinket-m0-circuitpython-arduino/circuitpython
3. Access the microcontroller CIRCUITPY drive following manufacturer instructions at https://learn.adafruit.com/welcome-to-circuitpython/the-circuitpy-drive
4. Delete the code.py file at the CIRCUITPY drive.
5. Rename the *Microcontroller_Main.py* file to *main.py* and upload it to the microcontroller CIRCUITPY drive.

### LI-6800 software configuration

1. From the GitHub repository, download the *LI-6800 Lamp Config*, *PAR_Ctrl.py*, *PAR_Ctrl_wFB.py* and *A-PPFD_Curve.py* files and copy them into a USB stick.
2. Insert the USB stick in the LI-6800 console and navigate to *Tools > Manage Files*.
3. Select the option *Copy files to LI-68000*.
4. Copy the *LI-6800 Lamp Config* file to the configuration directory */home/licor/configs*.
5. Copy the *PAR_Ctrl.py* and *PAR_Ctrl_wFB.py* files to the apps directory */home/licor/apps*.
6. Navigate to *Start Up > Configuration* and select the *Load configurations* option from the drop-down menu.
7. Select the configuration file you copied in step 4 and press the *Load* button.

## Appendix 2. Lamp PPFD output monitor and control calibration

This section provides step-by-step instructions to obtain the calibration coefficients needed to accurately monitor and control the lamp PPFD output. This section assumes that all building and configuration procedures defined in Appendix 1 are completed. The procedure is done in three steps. First, PPDF levels at various DAC1 control voltages are mesured. Then, a new photodiode multiplier is calculated. The new multiplier is used to monitor PPFD using the internal LI-6800-12A leaf chamber photodiode. Finally, the parameters of a linear model are calculated. These parameters are needed to predict PPFD output based on COB-LED heatsink temperature and DAC1 voltage. PPFD must be measured with an external device. Here we describe the procedures if using the Li-Cor Biosciences light meter (model LI-250A) equipped with the quantum sensor (model LI-190R Quantum); however, the procedure can easily be adapted if another model and/or brand is used instead.

### Taking needed measurements

1. Place the LI-6800 console, the leaf chamber, and the attached lamp in a dark room, preferably with air conditioning.
2. If possible, set the room temperature at a setpoint similar to the temperature at which the lamp will be used.
3. Remove the leaf chamber aperture insert from the bottom side of the LI-6800-12A leaf chamber.
4. Remove the leaf temperature sensor from the bottom side of the LI-6800-12A leaf chamber.
5. Install the 3 x 3 cm leaf chamber aperture insert only in the upper side of the LI-6800-12A leaf chamber.
6. Place the quantum sensor beneath the leaf chamber aperture insert in such a way that the sensing element is at the leaf plane. Use a laboratory stand and clamp to securely fix the sensor in place.
7. Increase DAC1 voltage gradually, from 0 to 5 V, in steps of 0.25 V. At each step, record *Qamb_in* and COB-LED heatsink temperature, using the LI-6800 console. At each step, record also the PPFD value measured by the quantum sensor installed in step 6, using an external light meter.
8. Repeat steps 5 to 7 using the 2 x 3 and 1 x 3 cm leaf chamber aperture inserts.
9. Repeat steps 5 to 8 at a different room temperature.
10. Prepare a data set in Excel with all measured data. Make sure to include the following variables as columns: DAC1 voltage, *Qamb_in*, PPFD measured by external sensor, COB-LED heatsink temperature and leaf chamber aperture insert area in cm^2^.

### Photodiode multiplier calibration

1. Obtain the original photodiode multiplier value from the leaf chamber memory. To do that, in the LI-6800 console, navigate to *Environment > Light > Ambient*.
2. Write down the original photodiode multiplier value and keep it.
3. From the measurements obtained during the previous subsection, filter out the data that has a saturated *Qamb_in* value. Use the remaining data in the following steps.
4. Calculate the slope of the linear regression between the measured PPFD using the external quantum sensor (X axes) and *Qamb_in* (Y axes).
5. Calculate the new photodiode multiplier by dividing the original multiplier value by the slope calculated in the previous step.
6. In the LI-6800 console, navigate to *Environment > Light > Ambient* and input the new photodiode multiplier.

### Lamp PPFD control model calibration

In this subsection, we provide detailed instructions to perform a multiple linear regression in SAS studio (other software can be used, for example, R or Python).

1. Create an account in SAS on demand for Academics at https://www.sas.com/en_us/software/on-demand-for-academics.html
2. Log in into SAS Studio.
3. Upload into SAS Studio the excel filet obtained in the first subsection of Appendix 2. Data file is loaded by navigating to *Server Files and Folders > Upload*.
4. Export the data to the *WORK* library folder as *IMPORT* library by navigating to *Tasks and Utilities > Utilities > Import Data*.
5. Start configuring a multiple linear regression task by navigating to *Task and Utilities > Tasks > Linear Models > Linear Regression*.
6. In the *DATA* tab, select the *WORK.IMPORT* library in the *Data* field.
7. Select the PPFD measured by the external quantum sensor as *Dependent variable*.
8. Select the DAC1, COB-LED heatsink temperature and aperture insert area as *Continuous variables*.
9. In the *MODEL* tab, click on the *Edit* button and add the model effects as outlined by Eqn. 1, without changing the estimates order. Make sure to deselect the use intercept option.
10. Run the task.
11. If all goes well, the r^2^ value for the model should be 1 or very close to 1, and the P-value for the model in the ANOVA table should be lower than 0.001.
12. Write down the four parameter estimates.
13. In the LI-6800 console, navigate to *Constants > User* and click on the *Edit/Reorder* option.
14. Select the user variable named Lamp_PAR_DAC1_Ctrl and click on the *Edit* button.
15. Replace the parameters named a1 to a4 with the parameters noted down in step 12. Make sure that the parameter order matches the order of both, Eqn. 1 and the model equation entered in step 9.
16. Save the changes and click on the *Edit/Reorder* option to deactivate the edition of User variables and constants.
17. Navigate to *Start up > Configuration* and select the *Save configurations* option form the drop-down menu.
18. Deselect all available settings to save, except for the user defined constants (*UsrDefCon*).
19. Save the changes to the configuration file.

## Appendix 3. Temperature control calibration

This section supplies step-by-step instructions to calibrate the control voltage over the TEMP_CTRL pin. This section assumes that all building and configuration procedures defined in Appendix 1 are completed. A multimeter is required for this calibration.

1. From the GitHub repository, download the *Zenner Diode Calibration_Data Template.xlsx* and *Curve_Fit_Zener_Diode.py* files. Place the files in the same directory.
2. Get ready the multimeter to measure the voltage between the TEMP_CTRL and ground pins at the microcontroller side. The point of measurement should be between the Zener diode and the microcontroller, and as close as possible to the microcontroller.
3. At the LI-6800 console, increase DAC2 voltage from 0 to 5 V in steps of 0.2 V. At each step, record DAC2 and TEMP_CTRL voltages in the excel file downloaded in step 1.
4. Execute the Python script downloaded in step 1.
5. Write down the estimated coefficients.
6. In the LI-6800 console, navigate to *Constants > User* and click on the *Edit/Reorder* option.
7. Select the user variable named Lamp_T_DAC2_V_Ctrl and click on the *Edit* button.
8. Replace the parameters named a1 to a4 with the parameters noted down in step 5. Make sure that the parameter order matches Eqn. 3.
9. Save the changes and click on the *Edit/Reorder* option to deactivate the edition of User variables and constants.
10. Navigate to *Start up > Configuration* and select the *Save configurations* option from the drop-down menu.
11. Deselect all available settings to save except for the user defined constants (*UsrDefCon*).
12. Save the changes to the configuration file.

## Appendix 4. Example uses

In this section, four lamp example uses are provided. This section assumes that all building and configuration procedures defined in Appendix 1 are completed.

### Direct voltage control of the lamp

Direct voltage control over the lamp is intended only for development and calibration procedures.

1. Navigate to *Start Up > Configuration* and select the *Load configurations* option from the drop-down menu.
2. Select the lamp configuration file and press the *Load* button.
3. Navigate to *Stat Up > Peripheral Setup > Console User I/O*.
4. Make sure the *Power5* supply is On, as it is needed to power the temperature sensor and microcontroller.
5. At the *DAC1* field enter the desired LED_CTRL voltage and press the *Done* button.
6. Navigate to *Measurements > Grid 4*.
7. Observe the user defined variables and constants related to the lamp. Notice that lamp PPFD changes (*Qin* variable) as the COB-LED heatsink temperature changes (*Lamp_T* user variable).

### Lamp PPFD control without feedback

1. Navigate to *Start Up > Configuration* and select the *Load configurations* option form the drop-down menu.
2. Select the lamp configuration file and press the *Load* button.
3. Navigate to *Programs > BP Builder > Open/New*.
4. Select the *PAR_Ctrl.py* file and press the *Start BP* button.
5. Navigate to *Constants > User*.
6. Input the desired PPFD in the *Lamp_PAR_Ctrl* user constant field.
7. Navigate to *Measurements > Grid 4*.
8. Observe the user defined variables and constants related to the lamp. Notice that the *Lamp_Qamb_in_Factor* user constant will not change, as the *PAR_Ctrl.py* script uses no feedback.
9. When you finish your experiment, navigate to *Programs > BP Monitor*.
10. From the table listing all BP running, select the *PAR_Ctrl.py* BP.
11. Press the *Cancel* button to stop the BP.

### Lamp PPFD control with feedback

1. Navigate to *Start Up > Configuration* and select the *Load configurations* option from the drop-down menu.
2. Select the lamp configuration file and press the *Load* button.
3. Navigate to *Programs > BP Builder > Open/New*.
4. Select the *PAR_Ctrl_wFB.py* file and press the *Start BP* button.
5. Navigate to *Constants > User*.
6. Input the desired PPFD in the *Lamp_PAR_Ctrl* user constant field.
7. Navigate to *Measurements > Grid 4*.
8. Observe the *Lamp_Qamb_in_Factor*, which indicates the level of feedback that the *PAR_Ctrl_wFB.py* script is applying.
9. When you finish your experiment, navigate to *Programs > BP Monitor*.
10. From the table listing all BP running, select the *PAR_Ctrl_wFB.py* BP.
11. Press the *Cancel* button to stop the BP.

### Manual A-PPFD curve

12. Navigate to *Start Up > Chamber Setup* and select the installed leaf chamber aperture insert.
13. Navigate to *Start Up > Configuration* and select the *Load configurations* option from the drop-down menu.
14. Select the lamp configuration file and press the *Load* button.
15. Navigate to *Start up > Warmup Tests*.
16. From the drop-down menu, select the option *Warmup Tests* and press the *Start* button.
17. If the warmup tests find no issues, continue to the next step; otherwise attend the issues before progressing.
18. Navigate to *Environment* tab and set the desired air flow, VPD, chamber CO_2_ concentration, fan speed, and leaf temperature.
19. Clamp the leaf to be measured.
20. Navigate to *Log Setup > Stability* and define the stability definition you wish to use (see section 4 of the LI-6800 manual for details).
21. Navigate to *Log Setup > Matching Options* and define the automatic matching schedule that you wish to use. It is recommended to match, at least, every 30 minutes and/or after a change in CO_2_ concentration (see section 4 of the LI-6800 manual for details).
22. Navigate to *Log Setup > Open a Log File* and create a new log file by pressing the *New File* button.
23. Navigate to *Programs > BP Builder > Open/New*.
24. Select the *PAR_Ctrl.py* file and press the *Start BP* button.
25. Navigate to *Constants > User*.
26. Input the desired PPFD in the *Lamp_PAR_Ctrl* user constant field. Within 10 s, the lamp should automatically adjust to the desired PPFD.
27. Navigate to *Measurements* tab and monitor the leaf response to the selected PPDF.
28. Wait for stability. When stability is reached, press the *Log* button.
29. Repeat steps 14 to 17 until you finish measuring all PPFD steps in the A-PPFD curve.
30. Navigate to *Log Setup > Logging to* ands press the *Close Log* button.
31. Navigate to *Programs > BP Monitor*.
32. From the table listing all BP running, select the *PAR_Ctrl.py* BP.
33. Press the *Cancel* button to stop the BP.

### Programmatic A-PPFD curve

1. Navigate to *Start Up > Chamber Setup* and select the installed leaf chamber aperture insert.
2. Navigate to *Start Up > Configuration* and select the *Load configurations* option from the drop-down menu.
3. Select the lamp configuration file and press the *Load* button.
4. Navigate to *Start up > Warmup Tests*.
5. From the drop-down menu, select the option *Warmup Tests* and press the *Start* button.
6. If the warmup tests find no issues, continue to the next step; otherwise attend the issues before progressing.
7. Navigate to *Programs > BP Builder > Open/New*.
8. Select the *A-PPFD_Curve.py* file and press the *Open BP* button.
9. Navigate to *Programs > BP Builder > Set*.
10. Expand the groups *Settings > Def. environmental settings*.
11. Adjust all environmental settings that the plant material/experiment needs. Here you define the flow, VPD, CO_2_ concentration, fan speed, leaf temperature, and chamber differential pressure to be set during the experiment. Notice that there are three variables for flow (*Low_flow_set, Medium_flow_set* and *High_Flow_set*); this is because the script uses different flow sets at different PPFD levels. The higher the PPFD, the higher the script sets the flow. This is to ensure maximum signal-to-noise ratio while keeping a ΔCO_2_ lower than 30 µmol·m^−2^·s^−1^ (as recommended by the equipment manufacturer). The PPFD thresholds defining each flow level are set in step 15.
12. Expand the groups *Settings > A-PPFD*.
13. Adjust the *PAR_levels* variable according to the plant material/experiment needs. The values you enter here are the PPFD values that will be used during the A-PPFD curve. The order in which are entered is the order in which the script will use them.
14. Adjust the *PAR_wait* variable according to the plant material/experiment needs. The values you enter here are the minimum waiting time (min) that script will keep each PPFD level. The order and number of items in the list must match the values entered in step 13.
15. Adjust the *Low_flow_PAR_end* and *High_flow_PAR_start* variables according to the plant material/experiment needs. These two variables are PPFD threshold levels defining three flow sets (see step 11 for explanation).
16. Press the *Save* button to skip all environment settings adjustments next time you use this script.
17. Clamp the leaf to be measured.
18. Navigate to *Log Setup > Stability* and define the stability definition you wish to use (see section 4 of the LI-6800 manual for details).
19. Navigate to *Log Setup > Matching Options* and define the desired automatic matching schedule. It is recommended to match, at least, every 30 min and/or after a change in CO_2_ concentration (see section 4 of the LI-6800 manual for details).
20. Navigate to *Log Setup > Open a Log File* and create a new log file by pressing the *New File* button.
21. Navigate to *Programs > BP Builder > Open/New*.
22. Select the script file you saved in step 16 and press the *Start BP* button.
23. Navigate to *Programs > BP Monitor* to monitor the two scripts that are running the A-PPFD curve. Notice that the main script automatically runs the *PAR_Ctrl.pu* script.
24. Navigate to *Measurements* tab and monitor the leaf responses to the PPDF levels.
25. When the script finishes the A-PPFD curve, navigate to *Log Setup > Logging to* ands press the *Close Log* button.
26. Navigate to *Programs > BP Monitor*.
27. From the table listing all BP running, select the *PAR_Ctrl.py* BP.
28. Press the *Cancel* button to stop the BP.

