## Supporting Information for "Open-source LED lamp for the LI-6800 photosynthesis system"

Vélez-Ramírez, A.I., *et al.*

### Supporting Information

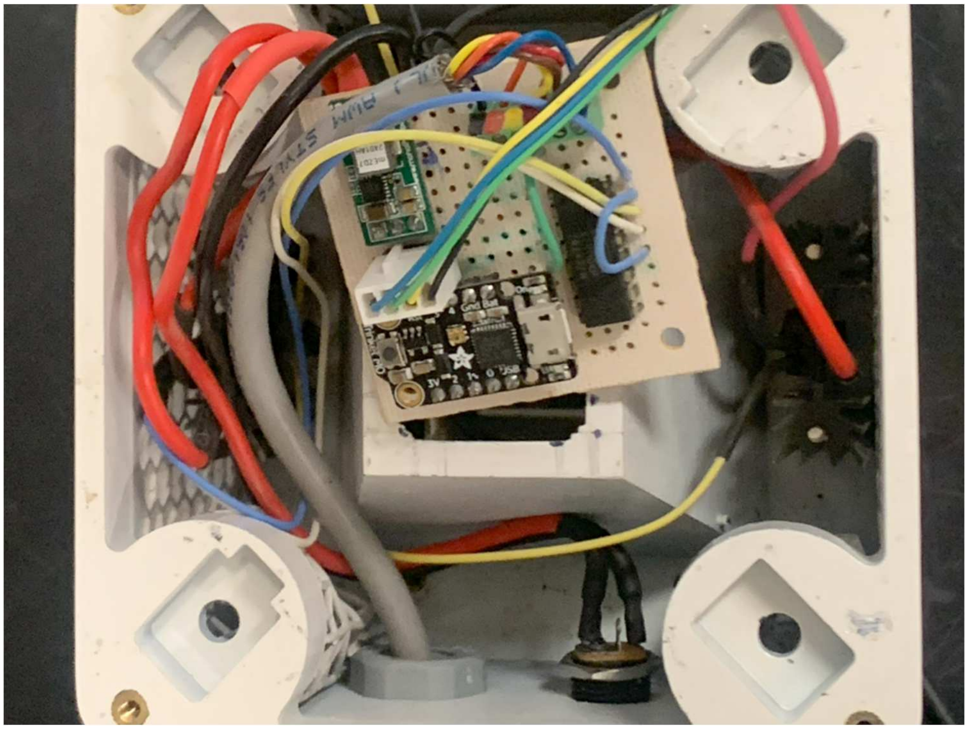

**Appendix S1.** Protoboard with the DC-DC voltage converter (top left), the XTR110 chip (Right) and.the microcontroller (Bottom left).

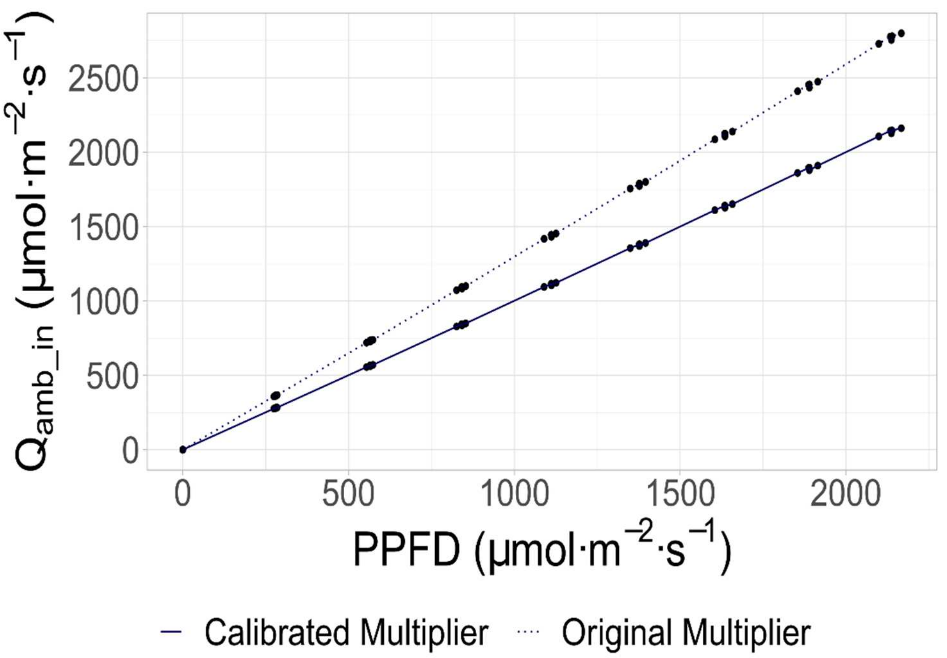


**Appendix S2.** Correlation between irradiance measured by an external quantum sensor (PPFD, X axes) and the internal photodiode ($Q_{amb\_in}$, Y axes). The dotted line indicates the correlation before multiplier calibration, and the continuous line depicts the calibration after the multiplier calibration.

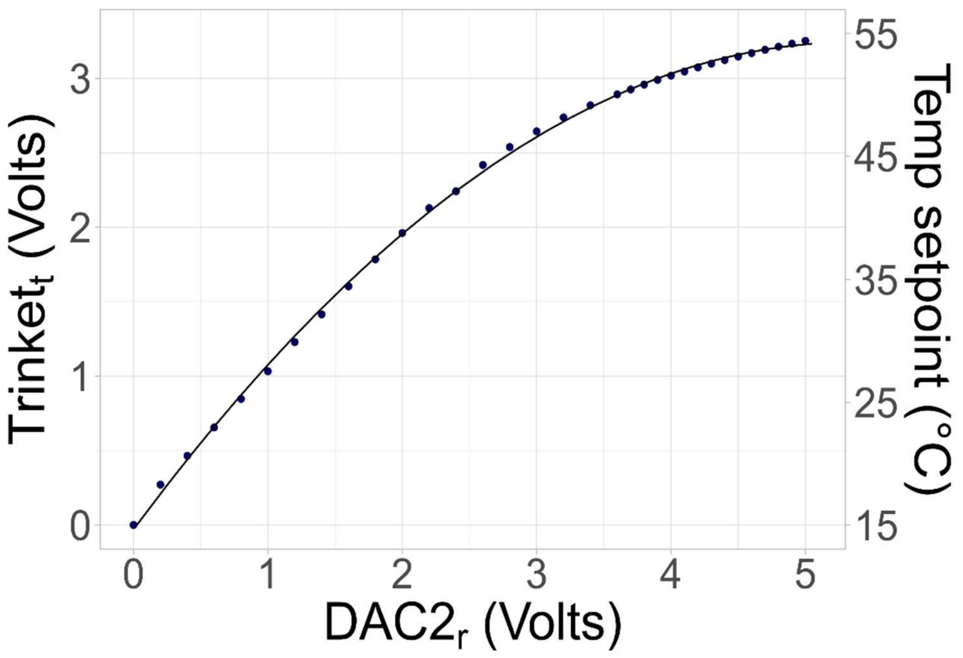


**Appendix S3**. Effect of the Zenner diode protection. The X axes depicts the required voltage at the LI-6800 *User I/O* DAC2 channel ($DAC2_r$) that is needed to achieve the target voltage at the Trinket M0 microcontroller side ($Trinket_t$, primary Y axes). Dots represent measurements and the line is a regression according to Eqn. 3. The second Y axes shows the resulting COB-LED heatsink temperature setpoint according to Eqn. 4.

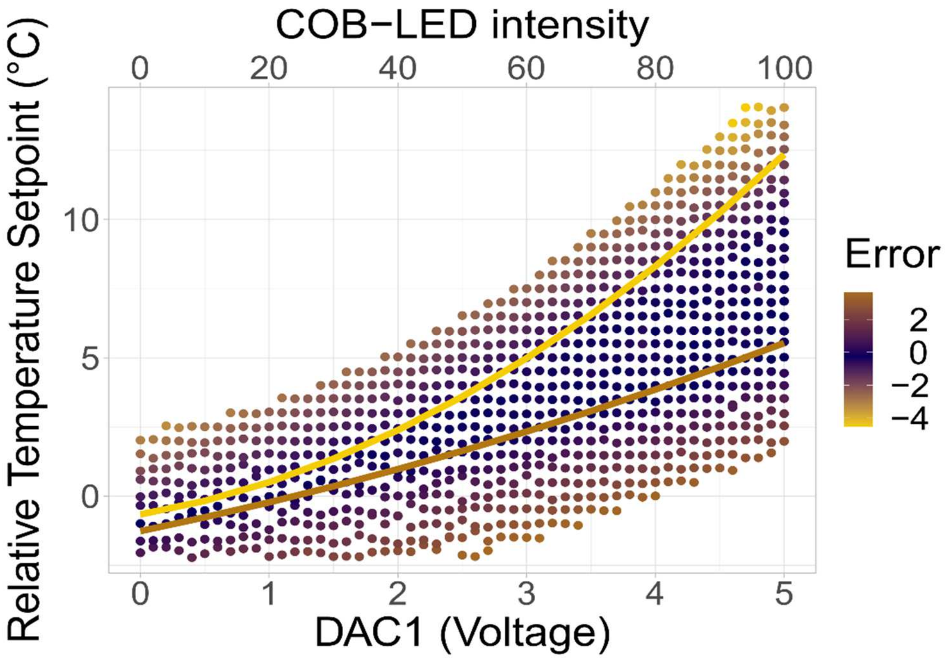


**Appendix S4**. Attainable COB-LED heatsink temperature setpoint. The X axes shows the voltage at the LI-6800 *User I/O* DAC1 channel, which controls LED current. That is, at a higher DAC1 voltage, the current flowing though the COB-LED, the emitted irradiance and the generated heat are larger. The Y axes depicts the COB-LED heatsink temperature setpoint, relative to air temperature. The dots depict 884 tested combinations of COB-LED intensity and heatsink temperature setpoint. The color scale indicates the deviation between the measured COB-LED heatsink temperature and its setpoint (Error in °C).

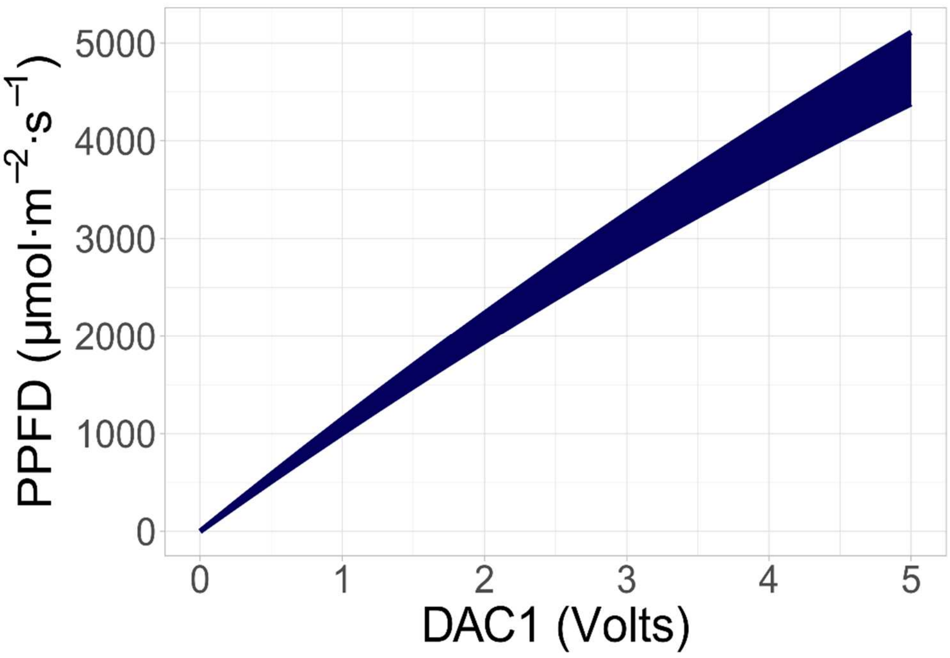


**Appendix S5.** Attainable PPFD. At any given DAC1 voltage, the shadow area indicates the attainable PPFD when heatsink is between 5 and 80°C.
